# Cell-surface glycans are quantitative reporters of Golgi dysfunction in single cells

**DOI:** 10.1101/2024.06.17.599374

**Authors:** Prachi Joshi, Aashish Satyajith, Debiprasad Panda, BR Rajeshwari, Mukund Thattai, Nagaraj Balasubramanian

## Abstract

Complex sugar polymers known as glycans contribute to the high molecular diversity of the eukaryotic cell surface. The types and levels of glycans on one cell can be sensed by other cells using carbohydrate-specific binding proteins such as lectins, enabling glycans to regulate cell-cell interactions. Glycans are covalently assembled onto proteins and lipids as they traverse the secretory pathway, a tightly regulated process known as glycosylation and carried out by Golgi-resident enzymes. Errors in glycosylation due to the dysfunctional trafficking of enzymes and substrates in the Golgi are implicated in human diseases. Here we ask how much information about Golgi dysfunction is encoded by surface glycans in single cells. This task is challenging due to high cell-to-cell variability in glycan levels. We exploited the loss-of-adhesion driven disorganization of the Golgi in mouse fibroblasts to generate a highly reproducible gradient of Golgi morphology phenotypes, by titrating the Arf1 inhibitor Brefeldin A (BFA). We measured the resulting distribution of cell-surface glycans in single cells using two fluorescently tagged lectin probes (ConA and WGA). A mathematical model of intracellular traffic, parameterized against measurements of Golgi fragmentation and endocytosis, quantitatively explains cell-surface lectin levels across time and BFA concentrations. We used this model to construct an optimal Bayesian decoder and showed that singlecell lectin measurements predict Golgi phenotypes with accuracy far greater than chance. By combining signals from two lectins we further improved prediction accuracy and speed. Such multi-lectin information may be exploited during natural cell-cell communication, and in the development of single-cell diagnostics.

---

The cell surface is an information-rich molecular tapestry. In eukaryotes, most cell-surface proteins and many lipids are covalently attached to glycans, a family of complex branched sugar polymers [1]. Glycans on one cell can interact with specific carbohydrate-binding proteins such as lectins on another cell [2], allowing these molecules to mediate cell-to-cell communication and inter-cellular interactions [3]. Cell-specific glycans regulate organismal development [4], while species-specific glycans prevent cross-species-fertilization [5], and facilitate self-nonself identification in immunity [6]. These examples demonstrate that glycans act as reporters of cellular identity or state. However, how such information is encoded and decoded at the single-cell level remains unclear. There is significant cell-to-cell variability in glycan types and levels, even among cells of the same developmental lineage [7]. How can glycans act as effective molecular reporters in the face of such variability?

Glycans are assembled onto protein or lipid substrates by glycosylation enzymes within the Golgi apparatus [8, 9]. The Golgi is organized as an ordered series of compartments, the cis-, medial- and trans-Golgi, each compartment containing a distinct complement of enzymes [10]. Proteins and lipids synthesized in the endoplasmic reticulum (ER) are trafficked through successive Golgi compartments on their way to the plasma membrane (PM). By constraining the order and duration in which growing glycans encounter glycosylation enzymes, Golgi compartmentalization regulates the spectrum of glycans made by a cell [11]. Disruption of Golgi organization and intra-Golgi trafficking impacts the types and levels of glycans on the cell surface [12–16]. Aberrant glycosylation is implicated in cancers [17, 18], pathogenic infections [19, 20], autoimmune disorders [19, 21] and genetic disease [22].

Here we ask whether cell-surface glycans can be used to probe Golgi dysfunction in single cells. This provides a concrete setting to explore how cell-to-cell variability constrains glycan information. We use a vesicle trafficking inhibitor to generate a spectrum of disrupted Golgi phenotypes, and measure the effect of this perturbation on cell-surface glycans. We incorporate these data into a mathematical model, and construct a decoder that reliably predicts the Golgi phenotype from single-cell surface glycan data. Our results highlight the potential of glycans as single-cell disease biomarkers [18], and also shed light on the mechanisms by which glycan information may be encoded and decoded in nature.

## Results

### Generating a gradient of Golgi fragmentation phenotypes

Golgi morphology can be perturbed by inhibiting the vesicle trafficking regulator Arf1 using Brefeldin A (BFA) [23–25]. We have previously shown that the application of BFA for varying times and concentrations generates a spectrum of Golgi morphology phenotypes in non-adherent mouse fibroblasts (WT-MEF cells) (Fig. 1A) [26, 27]. On loss of adhesion, most cells show a disorganized cis-Golgi morphology. The addition of BFA causes the cis-Golgi to become partially fragmented, and then completely fragmented as it falls back into the ER (Fig. 1B,C). The fraction of cells in each Golgi morphology class is extremely consistent at each BFA concentration and observation time.

**Fig. 1:**
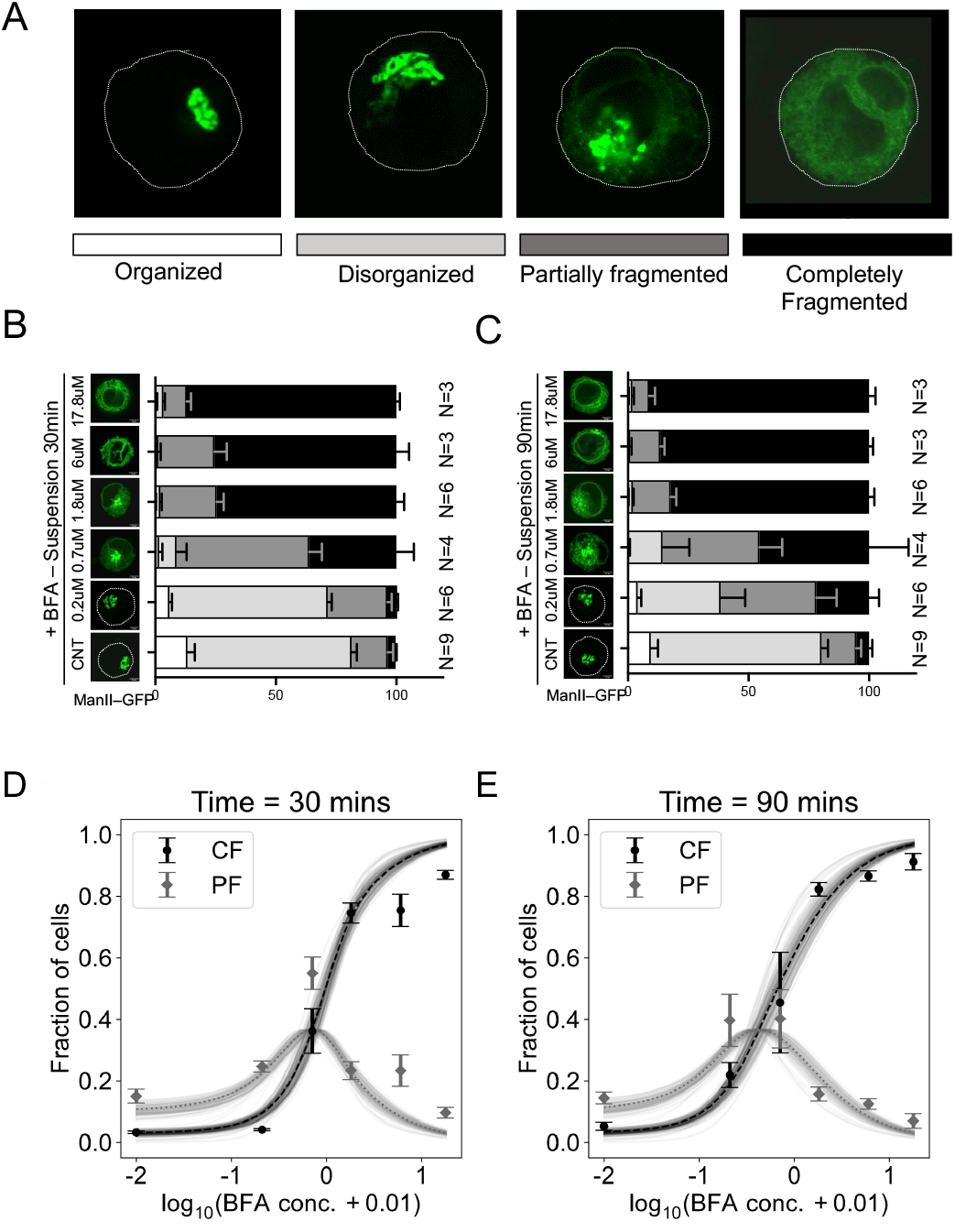
cis-Golgi fragmentation by Brefeldin A. We monitor the cis-Golgi in non-adherent WT-MEF cells using a fluorescently labeled cis-medial marker (Mannosidase II-GFP). (A) Representative images showing the four cis-Golgi morphology classes: organized, disorganized, partially fragmented, and completely fragmented. The completely fragmented class corresponds to the fallback of the entire cis-Golgi into the ER. (B,C) Distribution of cis-Golgi morphology in cell populations (n = 100) incubated for 30 min (B) or 90 min (C) with varying concentrations of BFA. Error bars show the SEM over N replicate populations. (D,E) The fraction of cells that are completely fragmented (black) or partially fragmented (grey) as a function of BFA concentration for 30min (D) or 90min (E). Data points and error bars show mean ± SEM of the observed values, taken from Fig. 1B,C. Dotted lines show the predictions of the 3-parameter cis-Golgi fragmentation model (Methods; Table 1). Light solid lines show variations in model predictions over 100 bootstrap replicates. The model is only fit to the completely-fragmented fraction, so the prediction to the partially-fragmented fraction is parameter-free.

To understand these dynamics, we developed a stochastic birth-death model for Golgi biosynthesis and fragmentation (Methods). We assume that the cis-Golgi is made up of discrete units, each of which is created by material from the ER [28, 29]. Arf1 inhibition reduces the flux of Golgi-to-ER COPI vesicles, leading to the accumulation of COPI-specific v-SNAREs in the cis-Golgi and promoting the fusion of cis-Golgi units with the ER. Under these assumptions the number of Golgi units is Poisson distributed [30] with a mean that depends on BFA concentration (c) and time (t):

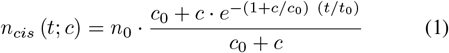

The fraction of cells with no cis-Golgi units is 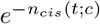, corresponding to the completely fragmented phenotype. Here, *n*_0_ = 3.6 is the average number of cis-Golgi units in steady state, in the absence of BFA; *c*_0_ = 0.15 µM is the BFA concentration at which the Golgi-to-ER COPI flux reaches its half-maximal level; and *t*_0_ = 83 min is the characteristic timescale on which cis-Golgi levels change (Table 1). These parameter values give a good qualitative fit to the observed fraction of cells in the completely fragmented class (Fig. 1D,E). The fraction of cells predicted to have a single cis-Golgi unit matches the data for the partially fragmented class with no further fitting.

**TABLE I:**
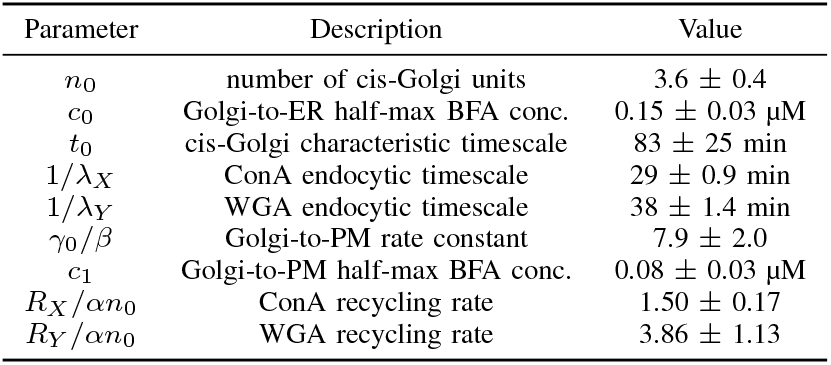
Parameter values (mean ± SD across bootstrap replicates)

| Parameter | Description | Value |
| --- | --- | --- |
| $n_0$ | number of cis-Golgi units | $3.6 \pm 0.4$ |
| $c_0$ | Golgi-to-ER half-max BFA conc. | $0.15 \pm 0.03 \mu\text{M}$ |
| $t_0$ | cis-Golgi characteristic timescale | $83 \pm 25 \text{ min}$ |
| $1/\lambda_X$ | ConA endocytic timescale | $29 \pm 0.9 \text{ min}$ |
| $1/\lambda_Y$ | WGA endocytic timescale | $38 \pm 1.4 \text{ min}$ |
| $\gamma_0/\beta$ | Golgi-to-PM rate constant | $7.9 \pm 2.0$ |
| $c_1$ | Golgi-to-PM half-max BFA conc. | $0.08 \pm 0.03 \mu\text{M}$ |
| $R_X/\alpha n_0$ | ConA recycling rate | $1.50 \pm 0.17$ |
| $R_Y/\alpha n_0$ | WGA recycling rate | $3.86 \pm 1.13$ |

### Measuring cell-surface glycans in single cells

We tracked the dynamics of cell-surface glycans in single cells using two lectins: ConA, which binds to oligo-mannose glycans, and WGA, which binds to glycans with sialic acid and N-GlcNAc terminal units [8]. These lectins were tagged with green and red fluorophores (Alexa488 and Alexa647). ConA-binding glycans are trafficked from the cis-Golgi to the trans-Golgi, where a fraction of them are further glycosylated to create WGA-binding glycans [12]. By probing glycans processed in different Golgi compartments, we can distinguish the effects of trafficking and glycosylation.

After confirming cell-surface lectin binding with confocal images (Fig. S1A), we standardized lectin labelling using flow cytometry. We opted for flow cytometry rather than imaging for quantification, taking advantage of the fact that our assay uses non-adherent cells. This allowed us to efficiently sample large cell populations and survey a large number of time points and BFA concentrations. We verified that single and dual-label assays, as well as fluorophore-swapped lectin constructs, produced consistent results (Fig S1), and that measured fluorescence intensities were linearly proportional to lectin concentrations (Fig. S2). Except for the endocytosis measurements described below, all subsequent experiments were done by flow cytometry using both fluorophore-swapped versions of the lectins as controls (CGWR, Con-Green/WGA-Red; CRWG, ConA-Red/WGA-Green). For clarity, in the main text we show results converted to CGWR fluorescence units.

Glycosylated proteins and lipids are endocytosed from the cell surface. Loss of adhesion and BFA treatment could both affect endocytosis [31–34]. Since we are interested in the impact of Golgi function on cell-surface glycans, we wanted to rule out any effect mediated by perturbations in endocytic rates. We measured the endocytosis of ConA-binding and WGA-binding glycans using a time-resolved assay (Fig. 2; Methods). Cell-surface lectin levels showed exponential decay, and the endocytic rate of ConA was consistently higher than that of WGA (Fig. 2B,C). The endocytic dynamics for each lectin were independent of BFA concentrations (Fig. S3).

**Fig. 2:**
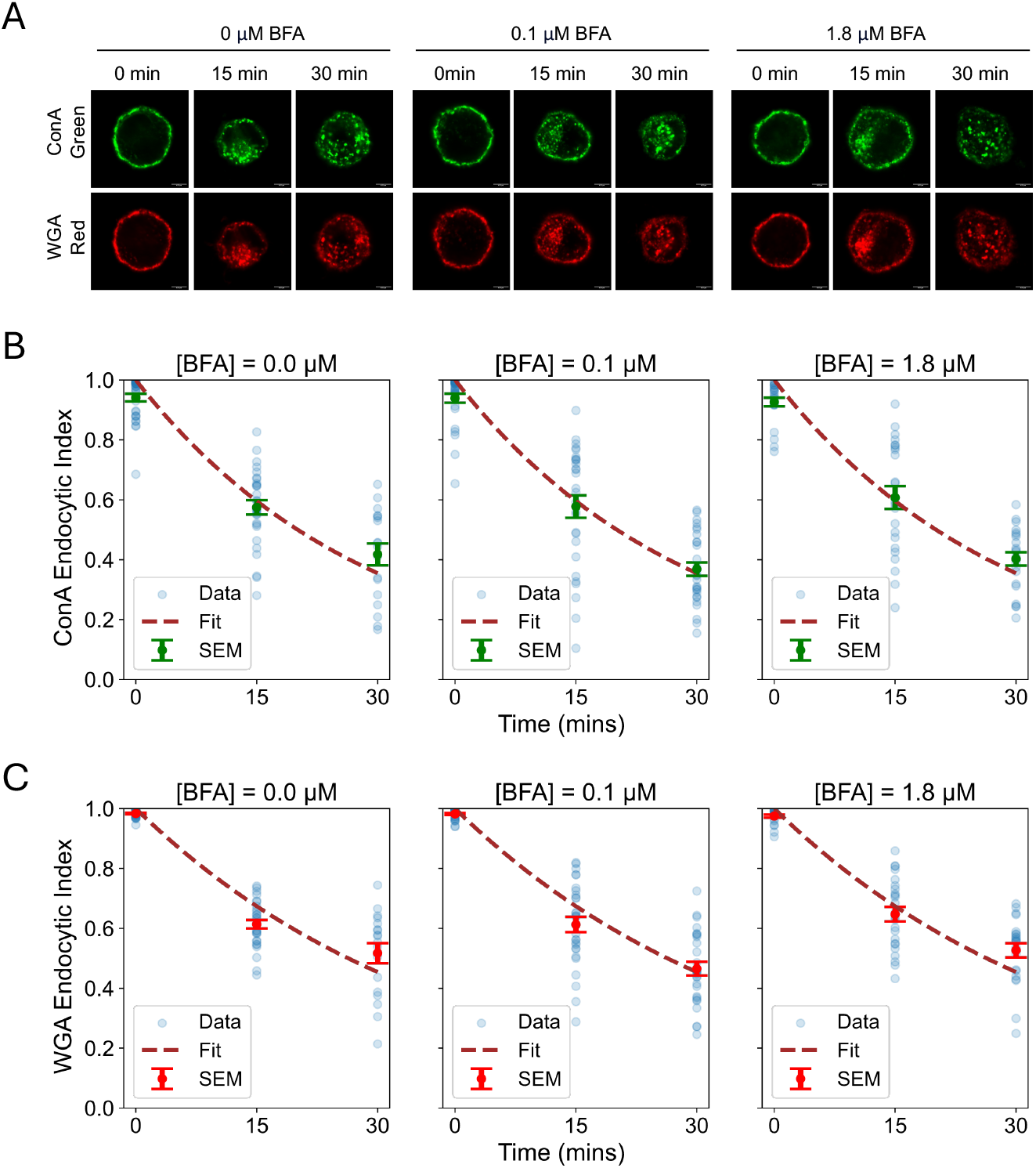
Endocytosis of cell-surface glycans. (A) WT-MEFs labeled with both ConA and WGA are fixed immediately, or maintained at 37°C for 15 min or 30 min before fixing. Confocal sections show representative images of lectin internalization in cells following each treatment. (B,C) We use a normalized endocytic index (EI; Methods) to measure the fraction of internalized lectins across time and BFA concentrations. Each time point shows data for *n*> 30 cells. Blue points show single-cell values, and green or red points show mean ± SEM. Dotted lines show the exponential fit to *EI* = *e* ^−*λt*^. Endocytic kinetics do not vary with BFA concentration (Fig. S3) but are different for ConA (B) and WGA (C): 1/λ_*X*_ = 29 min, 1/λ_*Y*_ = 38 min (Table 1).

### Modeling cell-surface glycan dynamics

Cell-surface glycan levels are determined by an interplay between synthesis and secretion from the Golgi, endocytosis, and recycling. We combined all these processes within a unified model of intracellular traffic and glycosylation (Fig. 3), summarized in the following equations:

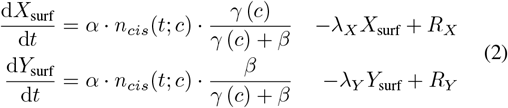

Here, the terms corresponding to cis-Golgi fragmentation (*n*_*cis*_(*t;c*)) and endocytosis (λ_*X*_, λ_*Y*_) have already been determined by our previous measurements. The parameters *α* and *β* can be absorbed by normalization. This leaves only four free parameters: two recycling rates *R*_*X*_ and *R*_*Y*_, and two parameters to define how BFA inhibits the rate constant of Golgi-to-PM transport: *γ(c)*= *γ*_0_/(*1 + c*/*c*_1_).

**Fig. 3:**
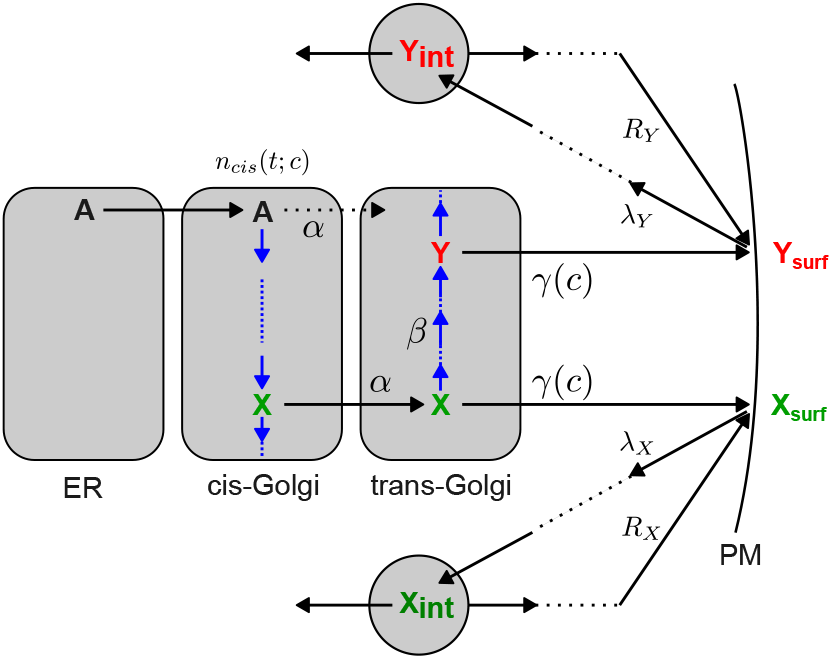
Modeling the synthesis and trafficking of glycans. For simplicity, we assume that the Golgi consists of two enzymatically distinct compartments, cis and trans. Oligo-mannose glycans (A) are synthesized in the ER and transported to the cis-Golgi, where they are further processed to form ConA-binding glycans (X). Glycans from the cis-Golgi are transported to the trans-Golgi with rate-constant *α*, so the flux of glycans emerging from the cis-Golgi is *α*_*cis*_ (*t;c*) (c represents the BFA concentration). X can be directly trafficked from the trans-Golgi to the plasma membrane with rate-constant *γ* (*c*), or it may be first glycosylated with rate-constant *β* to create WGA-binding glycans (Y). The fraction of X converted to Y is therefore *β*/(*γ* (*c*) + *β*). The trans-Golgi residence time is proportional to 1/*γ* (*c*). ConA- and WGA-binding glycans at the cell surface (X_surf_ and Y_surf_) are endocytosed at rates λ_*X*_ and λ_*Y*_. We assume that there are stable internal pools X_int_ and Y_int_ from which glycans are recycled to the plasma membrane at rates *R*_*X*_ and *R*_*Y*_.

This model is based on the following assumptions, supported by published data and our own observations. (1) The time taken for a glycan to go from the cis-Golgi to the plasma membrane is much shorter than the timescale over which the mean number of cis-Golgi units changes. This is justified since Golgi residence times range from 5 to 15 minutes [35, 36], whereas the observed cis-Golgi relaxation time is *t*_0_ ∼ 83 minutes (Fig. 1). (2) Endocytic rate constants are independent of time and BFA concentrations. This is consistent with our endocytosis measurements (Fig. 2). (3) Intracellular pools of endocytosed glycans are stable over the time course of the experiment, so the recycling rates are time-invariant [37]. (4) Glycosylation enzymes are not saturated [36], so enzymatic reactions obey first-order kinetics. (5) BFA does not directly affect enzymatic rates [38]. (6) BFA reduces the flux of glycans to the trans-Golgi via its influence on the number of cis-Golgi units [39] (Fig. 1). (7) BFA can inhibit the formation of clathrin-coated vesicles from the trans-Golgi to the plasma membrane, since these are regulated by Arf1 activity [39, 40].

### Impact of Golgi fragmentation on cell-surface glycan levels

The highly consistent effect of BFA on Golgi morphology in non-adherent cells provides a unique context to evaluate how Golgi perturbations affect Golgi-dependent glycosylation. We tracked cell-surface levels of ConA and WGA in single cells by flow cytometry, across the gradient of Golgi fragmentation phenotypes generated by BFA (Fig. S4; Methods). After scaling for cell size using forward scatter as a proxy, the empirical distribution of red and green fluorescence intensities were well-described by a bivariate log-normal distribution under each condition (Fig. S5). We henceforth use median fluorescence intensities as summary statistics for each population. To compensate for batch-to-batch variations in labelling efficiency, we normalized all fluorescence intensities by the intensity of the zero BFA control within each batch (Fig. S6). We refer to this normalized median value as the ConA or WGA signal. After this normalization, fluorescence values between fluorophore-swapped controls across all conditions showed excellent agreement (Fig. S7A), demonstrating the low measurement error of our assay.

At each BFA concentration, both ConA and WGA signals were seen to decrease from the 30 min to the 90 min time point, as expected when glycan synthesis is reduced (Fig. S7B,C). More interestingly, for fixed time, ConA signals showed a steep and consistent decrease as a function of BFA (Fig. 4A), while WGA signals showed a pronounced increase at low BFA (0.2 µM) followed by a subsequent decrease (Fig. 4B). There was a high degree of cell-to-cell variability within cell populations at each condition (Fig. S5). However, the pattern of variability (the orientation and scale of the empirical log-normal distributions) remained nearly constant across conditions (Figs. S8,S9). The coefficient of variation of ConA and WGA levels within each population was ∼ 0.3-0.4, comparable to the spread of ConA and WGA signals across conditions.

**Fig. 4:**
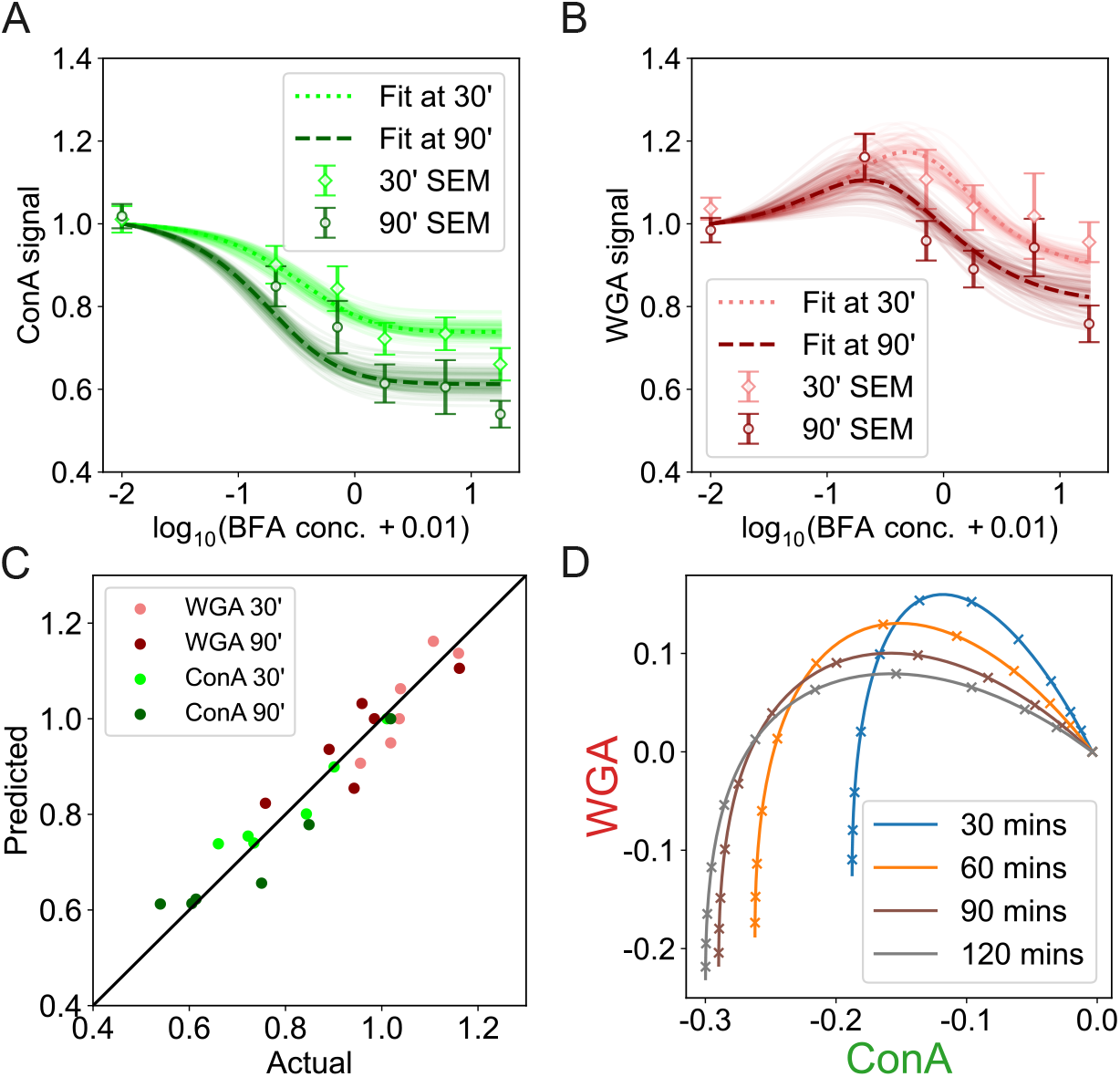
Cell-surface glycan dynamics can be explained by Golgi fragmentation dynamics. (A,B) Dynamics of cell-surface ConA (A) and WGA (B) signals, represented as a function of time and BFA concentration. The signal is defined as the median fluorescence intensity of a cell population, normalized by the intensity of the zero BFA control from the same batch (Methods). Data points and error bars show mean ± SEM of signals across replicate populations. The dotted lines show the predictions of the 4-parameter trafficking model. Light solid lines show variations in model predictions over 100 bootstrap replicates (Methods). (C) There is a strong agreement between predicted and actual values. (D) Predicted ConA and WGA signals as a function of time and BFA concentration in a 2D log-log plot (natural logs). The points marked ‘x’ correspond to a uniform grid of BFA concentrations in log space, used as a prior for decoding (Methods). This prior is chosen because it evenly samples the parabolic trajectories.

The observed ConA and WGA dynamics were well captured by the trafficking model Eq. 2, with just four free parameters (Fig. 4C). Most fitted parameter values fell within a tight uncertainty range (Table 1; Fig S10). The scale and timing of the observed variations are essentially fixed by *n*_*cis*_(*t; c*) and λ_*X*_, λ_*Y*_, showing that the overall decrease in ConA and WGA signals is driven by reduced flux from the cis-Golgi and ongoing endocytosis. The recycling rates *R*_*X*_ and *R*_*Y*_ set the baseline level of surface glycans at high BFA concentration, where synthesis is expected to be negligible (note that recycling may be overestimated due to background fluorescence). The remaining two parameters, specifying the BFA-dependence of the Golgi-to-PM transport rate *γ*(*c*) = *γ*_0_/(1 + *c*/c_1_), are required to capture the “bump” in WGA signals. This suggests that BFA addition increases the trans-Golgi residence time, amplifying the conversion of ConA-binding glycans to WGA-binding glycans. However, this effect would need to be checked by independent measurements.

### Using glycans to probe Golgi fragmentation in single cells

According to our birth-death model, (Eq. 4) cis-Golgi units are gained or lost every *t*_0_/(2*n*_0_) ∼10 min; in comparison, cell-surface glycans have a lifetime of over 1/λ_*X*_ = 30 min before being endocytosed (Table 1). Surface glycan levels in a single cell therefore reflect the average number of functional cis-Golgi units in which these glycans were processed over the past 30 minutes. This explains why lectin signals form a unimodal distribution (Fig. S5): at a given BFA concentration, all cells in a population have similar time-averaged cis-Golgi levels, though they may have very different instantaneous cis-Golgi morphologies. Suppose cells could have been drawn from one of several populations, each characterized by a distinct value of n_*cis*_ (Fig. 5A). By measuring lectin levels on a single cell, can we infer which population it belongs to?

**Fig. 5:**
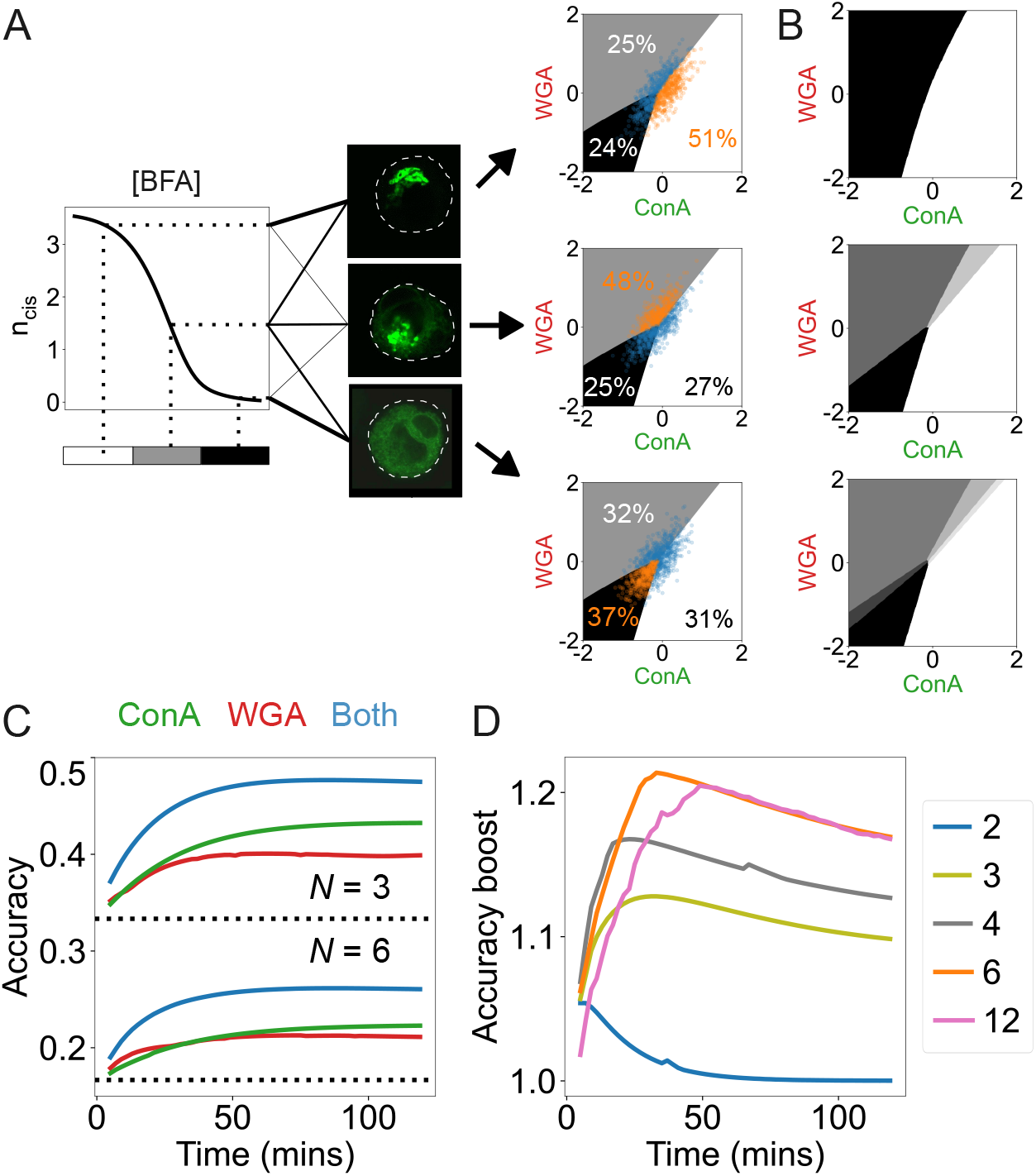
Decoding Golgi phenotypes from single-cell measurements. (A) Left: We assume a logarithmic prior of 12 BFA concentrations (Fig. 4D) and split these into *N* equiprobable ranges (grey shading). Populations at distinct BFA concentrations contain distinct mixtures of Golgi morphologies (illustrated by lines of varying thickness). We pool all populations corresponding to a range of BFA concentrations into *N* = 3 meta-populations (white, grey and black). Right: The *N* = 3 decoder for the 30 min time point, showing the regions of fluorescence space that map to each meta-population. The scatterplots show synthetic datasets of 1000 cells generated according to the predicted meta-population distributions. Percentage values show the fraction of cells that lie in each region. Orange points show the cells that decode to the correct meta-population. Note that the decoders do not predict the Golgi morphology of a single cell, they predict that the cell was drawn from a meta-population with a specific mixture of Golgi morphologies. (B) Decoders for *N* = 2, 4, 6. All decoders are shown in the log-log space of ConA and WGA signals (natural logs). (C) Accuracy of decoding (averaged across *N* meta-populations) as a function of time, using only ConA, only WGA or both lectins together. Black dotted lines indicate the accuracy 1/ *N* expected by chance. (D) The boost in accuracy from using two lectins, compared to using only ConA, for various values of *N*. (The small spikes seen in panels (C) and (D) are an artefact of the discrete concentration prior.)

The optimal Bayesian decoder provides the best answer to this question, in the sense that it minimizes the probability of error [41] (Methods). To set up the decoder, we start with an ordered series of cell populations, with decreasing levels of *n*_*cis*_. These are generated according to predicted means and covariances for 12 BFA concentrations, uniformly distributed in log space (these concentrations are sufficient to capture the full range of ConA and WGA variation, see Fig. 4D). We pool these into *N* meta-populations, each corresponding to a contiguous range of BFA concentrations. *N* sets the resolution of the inference task (Fig. 5A, left); we must construct different decoders for each choice of time and resolution (Fig S11). The decoder is implemented as a gating operation on single-cell fluorescence data: it splits the space of fluorescence values into *N* regions corresponding to different meta-populations. If a single cell falls into a certain region, our best guess is that it was drawn from the corresponding meta-population. In Fig. 5A,B we show dual lectin decoders for the 30 min time point, for *N* = 2, 3, 4, 6; these decoders have a non-trivial polar structure due to the parabolic trajectory of the means (Fig. 4D). Note that the decoder resolution is not related to the number of Golgi morphology classes, which is always four (Fig. 1A); the resolution is the number of meta-populations, each corresponding to a distinct mixture of Golgi morphologies. Interestingly, the decoders typically split into three large regions and several smaller regions, suggesting that the *N* = 3 decoder is the natural choice for this specific task. This means we can efficiently decode to three meta-populations, corresponding to high, intermediate, and low levels of *n*_*cis*_.

We can determine decoder accuracy by repeating the inference task over many cells (Fig 5A, right). We find that the decoders predict the correct meta-population far better than chance (Fig 5C). Decoders that integrate information from two lectins perform significantly better than those that use a single lectin (Fig 5C). The performance of the decoders tends to increase with time since BFA addition. Dual-lectin decoders perform better than single-lectin decoders, with a boost in accuracy around the 30 min point (Fig 5D), reminiscent of dynamical information synergy [42]. Using two lectins thus alleviates the speed-accuracy tradeoff: for *N* = 3, the accuracy achieved by a single-lectin decoder at 50 min is matched by that of a dual-lectin decoder at 15 min.

## Discussion

How do changes in the organization of the Golgi apparatus impact its function? The question is surprisingly difficult to address, due to the complex and dynamic nature of this organelle. Drastic perturbations, for example, the mislocalization of Golgi-resident enzymes, can change both the type and level of glycans made by a cell, which can be easily discerned at the population level [14–16]. Here, we are interested in the effect of more subtle Golgi perturbations and how they manifest at the scale of a single cell. We have two central results. First, subtle changes to the Golgi cause proportionate, quantitative changes in glycan levels (Fig. 4). Second, by using a mathematical model to invert this link, we can effectively infer Golgi morphology by measuring cell-surface glycans in single cells (Fig 5).

Our results can be framed as a signal-to-noise question: can we detect changes in the median phenotype of a cell population under varying conditions, when there is large variability between cells within each population? To generalize our results, it would be important to identify the drivers of cell-to-cell variation. The observed differences in Golgi fragmentation across cells (Fig. 1) would produce correlated changes in the levels of all glycans. Other sources of variation, for example in enzymatic and recycling rates, would produce uncorrelated stochastic effects across glycans. Heterogeneity in Golgi mor-phologies and glycosylation signatures may influence glycan-dependent processes in development and disease [43–45]. Methods to reliably measure Golgi function and dysfunction in single cells could in principle be used to develop disease biomarkers [18, 46]. Cancer cells with distinctly altered Golgi organisation [47–49] and glycosylation signatures [50–52] could be direct candidates to test this hypothesis. However, the application of our decoders in single-cell diagnostics would first require a reduction in measurement noise, since the pre-processing we do to reduce batch-to-batch variation depends on population-level data (Fig. S6).

Mathematical models play two distinct roles in our analysis. At one level they are merely curve-fitting tools that allow us to interpolate between and compress a large amount of experimental data into a few parameter values. Our results on decoder accuracy only depend on the analytic curves being good representations of the data, regardless of the ontological content of the underlying models. At another level, the fact that we can parsimoniously explain the observations using just a handful of parameters (Table 1) suggests that the models do capture aspects of the underlying cell-biological processes. The stochastic model of cis-Golgi biosynthesis provides a mechanistic basis for the BFA-driven fallback of the cis-Golgi into the ER (Fig. 1). The effect of BFA on the trans-Golgi is more subtle. Upon loss of adhesion, the trans-Golgi becomes dispersed, with a loss of active Arf1 and GBF1 [27]; but BFA does not further perturb its morphology (Fig. S12). Our trafficking model suggests that glycosylation continues to occur in the trans-Golgi following BFA addition, but the increased compartmental residence time changes the proportion of different glycan types that are synthesized (Fig. 3).

More work would be needed to understand how glycans mediate information transfer between cells in natural settings. We have shown how cell-to-cell variability in glycan synthesis constrains the amount of information that can be encoded by a “sender” cell (Fig 5). A complementary study has shown how noise in lectin signaling limits the fidelity with which glycans levels can be decoded by a “receiver” cell [2, 53]. The use of multiple glycans boosts the available information (Fig 5D) at the cost of requiring a more complex decoding system. The differential responses of ConA and WGA signals to the same BFA-driven perturbation are central to the performance of our dual-lectin decoder (Fig. 4D). This illustrates a general principle: more information can be gained by combining diverse glycans [54]. Better decoder accuracy depends not just on the number of distinct glycans, but rather on the choice of relevant pairs or triplets of glycans processed in distinct ways across Golgi compartments. The Golgi apparatus can potentially synthesize an astronomically large number of glycan variants [12]. It is therefore likely that glycan diversity is an essential feature of cell-cell communication, and that cells are able to incorporate measurements of multiple glycans in sophisticated ways. When it comes to understanding how glycan information is encoded and decoded, we have only scratched the surface.

## Materials and Methods

### Reagents

Accutase (Cat. No. #A6964), Brefeldin A (Cat. No. #B7651) and DMSO (Cat. No. # D2438) were purchased from Sigma. Lipofectamine 2000 (Cat. No. # 11668019) was purchased from Invitrogen. Fluorophore conjugated lectin probes were purchased from Molecular Probes - ConA-Alexa488 (Cat. No. #C11252), ConA-Alexa647 (Cat. No. # C21421), WGA-Alexa488 (Cat. No. #W11261), WGA-Alexa647 (Cat. No. #W32466). Fluoramount-G (Cat. No. #0100-01) was purchased from Southern Biotech. The GalTase-RFP and Mannosidase II-GFP construct was obtained from Dr. Jennifer Lippincott-Schwartz.

### Cell culture and transfections

Wild-type mouse Embryonic Fibroblast cells (WT-MEFs) were obtained from Dr. Richard Anderson (University of Texas Health Sciences Centre, Dallas, TX). Cells were cultured using Gibco DMEM from Thermo Fisher Scientific, with the addition of 10% Penstrep and 5% FBS. 0.05% Trypsin Accutase (for lectin labelling experiments) was used for detaching cells, and excess culturing medium was used for neutralizing the action of Trypsin. For transfection studies, cells were seeded in 6 cm dishes to attain a confluency of 60% and allowed to attach and spread for 4 hours. Using Gibco OptiMEM medium and transfection agent Lipofectamine 2000 from Thermo Fisher Scientific, a transfection mix was prepared with 3 µg of Mannosidase II-GFP. The mixture was kept at room temperature for 30 minutes before being added to the cells seeded in 6 cm dishes. Media in transfected dishes was changed 12 hours post-transfection, and cells were used for experiments 36 hours post-transfection.

### Suspension of cells treated with BFA

Cells in culture were grown up to 70% confluency in 10 cm or 6 cm dishes. Cells were serum deprived using low serum DMEM (0.2% FBS) for 14.5 hrs before detaching cells for the suspension assay. Cells were detached using 0.05% Trypsin or Accutase (for all lectin labelling experiments), washed with low-serum medium and processed further. One aliquot was kept aside from collected cells and processed at the 5 min (just detached) timepoint when required. Cells were gently mixed with a low-serum medium containing 1% methylcellulose for the suspension assay. Increasing cell numbers accompany a comparable increase in suspension volume. DMSO or BFA was added to this suspension at a range of concentrations (0.2 µM, 0.7 µM, 1.8 µM, 6 µM and 17.8 µM) as required for the experiment. The suspension tube was gently mixed to ensure DMSO/BFA was distributed uniformly. This cell suspension mix was incubated at 37°C with 5% CO2 for the required time. Methylcellulose was diluted and washed using a cold, low-serum medium post-suspension. Cells were eventually fixed or labelled with lectin as needed. Fixed samples were mounted on slides using Fluoramount-G, dried and then used for confocal imaging. This procedure was followed for cells transfected with Mannosidase II-GFP when required.

### Confocal imaging for classification of Golgi phenotypes

Fluoramount-G mounted slides were imaged on a Zeiss confocal (LSM710) using a 63X oil immersion objective. Cross-sectional images were acquired and captured with averaging of 4, a scan speed of 5 and a zoom of 5 for non-adherent cells. The equatorial plane was shot for each cell to capture images of cell surface lectin binding, identified by the largest perimeter. For profiling cells, a minimum of 100 and a maximum of 200 cells were counted and manually categorized according to their Golgi organization phenotype: (i)organized Golgi, compact; (ii) disorganized Golgi, opened up; (iii) partially fragmented, opened up with some fallback to the ER; (iv) completely fragmented, complete fallback to the ER [27].

### Birth-death model for Golgi biosynthesis and fragmentation

COPI vesicles mediate retrograde transport from the cis-Golgi, and require specific v-SNAREs to drive their fusion to the ER [55]. We assume the per-membrane-area concentration of these v-SNAREs on the cis-Golgi obeys

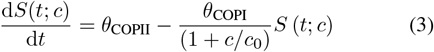

where the first term corresponds to the arrival of v-SNAREs from the ER on COPII vesicles, and the second to their removal from the cis-Golgi by COPI vesicles. The rate of COPI vesicle formation is inhibited by increasing BFA concentrations (c). We assume v-SNAREs rapidly equilibrate, reaching a quasi-steady-state level *S*(*c*)= (*θ* _COPII_/*θ*_COPI_) (1 + *c*/*c*_0_).

We model the cis-Golgi as being made up of discrete units [28, 29]. These units are created by material from the ER, and they can fuse back to the ER at a rate proportional to the v-SNARE concentration. The expected number of cis-Golgi units is therefore given by:

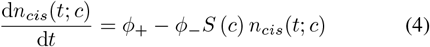

Since birth and death are stochastic, there will be fluctuations around this expected value. It can be shown that the number of cis-Golgi units under this stochastic birth-death process is given by a time-dependent Poisson distribution [30] whose mean is specified by integrating Eq. 4 to give Eq. 1 of the main text, with parameters *t*_0_ = (*θ*_COPI_/*θ*_COPII_) /*ϕ*_—_, and n_0_ = *ϕ*_+_*t*_0_. Since *n*_*cis*_ (*t;c*) is Poisson distributed, the fraction of cells with zero Golgi units is given by 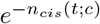. We fit this prediction against the observed fraction of cells with completely fragmented cis-Golgi.

### Lectin labelling and flow cytometry analysis

Cells detached using Accutase and held in suspension as described above were given a PBS wash to remove residual media and then reconstituted in a defined volume of PBS for further lectin labelling. A range of lectin concentrations was evaluated to optimise the concentration of lectin (ConA, WGA) tagged with a fluorophore (Alexa488, Alexa647) to be used for these studies (Fig. S3). When analysed by flow cytometry, a lectin concentration that gave intensity measurements in the range of 5000 – 10000 arbitrary units was selected. Each lectin-fluorophore combination was tested individually before optimized concentrations were combined for dual labelling. The final concentrations were: ConA-Alexa488, 50 ng/µL; ConA-Alexa647, 5 ng/µL; WGA-Alexa488, 0.5 ng/µL; WGA-Alexa647, 0.25 ng/µL.

For dual labelling, cells were incubated with optimal concentrations for ConA and WGA tagged with different fluorophores (Alexa488 vs. Alexa647). Each labelling was done with both fluorophore-swapped lectin combinations (ConA-Alexa488 + WGA-Alexa647; ConA-Alexa647 + WGA-Alexa488). The labelling reaction was kept on ice in the dark for 15 minutes, followed by two washes with cold PBS. For flow cytometry analysis, PFA (3.5%) was used to fix the lectin-labelled cell samples resuspended in cold PBS (350 µL). Samples were run on BD Celesta Flow Cytometer. A morphologically uniform cell population defined using their forward and side scatter (FSC and SSC) plot was selected using polygon gating for data acquisition. More than 5000 individual events were recorded in the gated area for every sample. This was further analysed using FlowJo software to get single-cell data for each sample.

### Testing endocytosis of cell surface lectins

Cells detached using Accutase and held in suspension as described above (with DMSO/BFA - 0.1 µM and 1.8 µM) were given a PBS wash to remove residual media and then reconstituted in a defined volume of PBS for further lectin labelling. This labelling was done as described above. Labelled cells were equally distributed for three time points – 0 min, 15 min and 30 min of endocytosis. Cells were kept in 500 µL low-serum media (with DMSO/BFA-0.1 µM and 1.8 µM) at 37°C to allow endocytosis of surface-labelled lectins at respective time points. These cells were washed with cold PBS, spun down and fixed with 3.5% PFA. All samples were mounted on slides using Fluoramount-G, allowed to dry and then used for confocal imaging. Cell surface lectin labelled control and BFA-treated live cells were (i) fixed immediately as a 0 min pre-endocytic control or (ii) incubated at 37°C for 15 min or 30 min to allow for lectin endocytosis. Using confocal cross-section images, cells fixed at 0 min showed an exclusively membrane localized lectin binding, that is visibly endocytosed into the cell after 15 and 30 min of incubation.

For quantification, the Python module Cellpose [56] was used to identify the cell boundary. Computed boundaries were manually inspected and found to be well determined for 234 out of 270 cells; these were retained for further analysis. The saved boundaries were imported into Fiji’s image analysis software as .roi files [57]. The cell periphery was defined as the region obtained by scaling the original boundary from 85% to 105%, and the interior was defined as the region within the 85% limit. We computed an endocytic index E for each cell and labelled lectin, defined as the fraction of pixels in the periphery whose intensity is greater than the median intensity of the interior of the cell. This index is robust to changes in background fluorescence and overall fluorescence scales. We found that BFA had no effect on observed endocytic indices (Fig. S3). The endocytic indices of ConA and WGA, pooled across all BFA concentrations, were fit to first-order exponential decay by least-squares minimization: 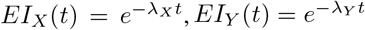.

### Single-cell lectin data normalization

(i) Red and green intensities of single cells were consistently proportional to their forward scatter (FSC) in the flow-cytometric measurements (Fig. S5A,B). We interpreted this as a cell size effect, and corrected for it by scaling all intensities by the FSC. (ii) Scaled red and green intensities of cell populations showed an approximate bivariate log-normal distribution, across conditions (Fig. S5C). We used robust estimation, implemented in the Python module astroML [58], to determine the distribution means and covariances (Figs. S8,S We then checked if 95% of the cell population fell within the 6*σ*-contour of the fitted distribution. Of the 237 flow cytometry experiments performed (across time, BFA concentrations and replicates) 207 satisfied this condition and were retained for further analysis. (iii) Sample preparation was done in batches, as shown in Fig. S4. There were large differences in absolute fluorescence levels across batches (Fig. S6, left column). To compensate for this, we scaled all fluorescence signals for each batch by the geometric mean of the 5 min, 30 min and 90 min 0 µM BFA controls in the same batch. Fluorescence values were tightly distributed following this normalization (Fig. S6, right column). We refer to this final normalized intensity as the red or green signal.

### Fitting trafficking model to lectin data

For every condition (combination of BFA and time) there were at least nine biological replicates, and at least 20 replicates for the 0 BFA controls. We computed the geometric mean of red signals, and the geometric mean of green signals, across replicates for each condition and fluorophore configuration. We then compared the mean red and green signal across equivalent experiments (i.e. red of CRWG against green of CGWR; or red of CGWR against green of CRWG). These showed a tight linear relationship (Fig. S7A). We therefore pooled all the data across both fluorophore-swapped controls for subsequent analysis, using the best fit line from Fig. S7A to convert CRWG measurements into CGWR units. We solved Eq. 2 with initial conditions *X*_surf_ = 1 and *Y*_surf_ = 1, to enable a direct comparison with normalized lectin signals. We inferred model parameters by least-squares fitting, using Python’s SciPy module [59]. To estimate confidence intervals for each parameter, we repeated the fitting procedure over 100 bootstrap replicate datasets. These datasets were synthesized by randomly sampling the ConA and WGA signals with replacement for each time and BFA concentration (except the 0 µM BFA controls which were earlier used to normalize intensities).

### Decoder analysis

We have determined how the median ConA and WGA signals vary with time and BFA concentrations. To build a decoder we also needed to model covariances. The elliptical 10-contour in log-log space provides a geometric summary of the covariance of a bivariate log-normal distribution. We found that the shape of this ellipse (its angle, scale and aspect ratio) were essentially independent of time and BFA concentration for the CGWR and CRWG constructs (Fig. S8, S9). We therefore modeled the covariance as constant, averaging the angle, major axis and minor axis across conditions. Since the red and green signals were converted to CGWR units for fitting the trafficking model, we used the average CGWR covariance for the decoder analysis. From the behavior of the medians and the covariances, we know the population distribution *p* (*g, r*|*c, t*). We construct decoders in log-log space, and *g* and *r* represent the natural logs of the ConA and WGA signals.

We assume the BFA concentration is chosen from a uniform prior across 12 values in µM: 0.0, 0.02, 0.04, 0.08, 0.16, 0.32, 0.64, 1.28, 2.56, 5.12, 10.24, 32.0. We split these into *N* equiprobable ranges of size 12/*N*, for *N* = 2, 3, 4, 6, 12. We then superpose distributions for all the concentrations in each range, to generate *N* meta-population distributions. The probability that meta-population *B* results in ConA signal *g* and WGA signal *r* is *p*_*B*_(*g, r*|*t*) = Σ_*c* ∈*B*_ *p* (*g, r*|*c, t*). For optimal Bayesian decoding we must find the most likely meta-population that generated a given *g, r* signal:

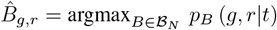

where ℬ_*N*_ denotes the set of *N* meta-populations. The decoder splits the fluorescence space into distinct regions, each mapping to distinct most-likely meta-populations (Fig. 5).

The accuracy of a decoder for meta-population *B* at time *t* is 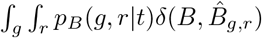 *dr dg* where 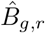 is the inferred meta-population at signal *g, r*, and 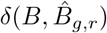 returns 1 if its inputs match, 0 otherwise. For evaluating this integral numerically, we considered the range of g and r from −2 to 2, which covered more than 99.999% of the signal probability distribution *p* (*g, r*). The overall accuracy of the decoder is

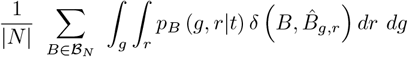

For decoding using a single glycan we proceed exactly as described above, replacing the joint distribution *p* (*g, r*|*c, t*) with the marginal distribution *p* (*g*|*c, t*) or *p* (*r*|*c, t*).

### Code availability

All the code and data used are available online at https://gitlab.com/aashishsatya/decoding-golgi-perturbations.

## Acknowledgements

MT acknowledges support from the Simons Foundation (287975). MT thanks Kaadambari and Raghav for their inputs on the figures. AS would like to thank Anton Swaminathan Iyer for pointing him to the Cellpose module, and his colleagues at Simons Centre, NCBS for helpful discussions.

NB acknowledges support from SERB (CRG/2022/001813) and IISER,Pune. PJ would like to thank IISER, Pune, for fellowship support. We acknowledge support from the IISER,Pune microscopy and flow cytometry facilities.

## Author Contributions

NB and MT conceived the project. PJ, DP and BR carried out the experiments. MT and AS developed the models. AS and PJ carried out the analysis. All authors wrote the paper.

