## Supplementary Figures for "Cell-surface glycans are quantitative reporters of Golgi dysfunction in single cells"

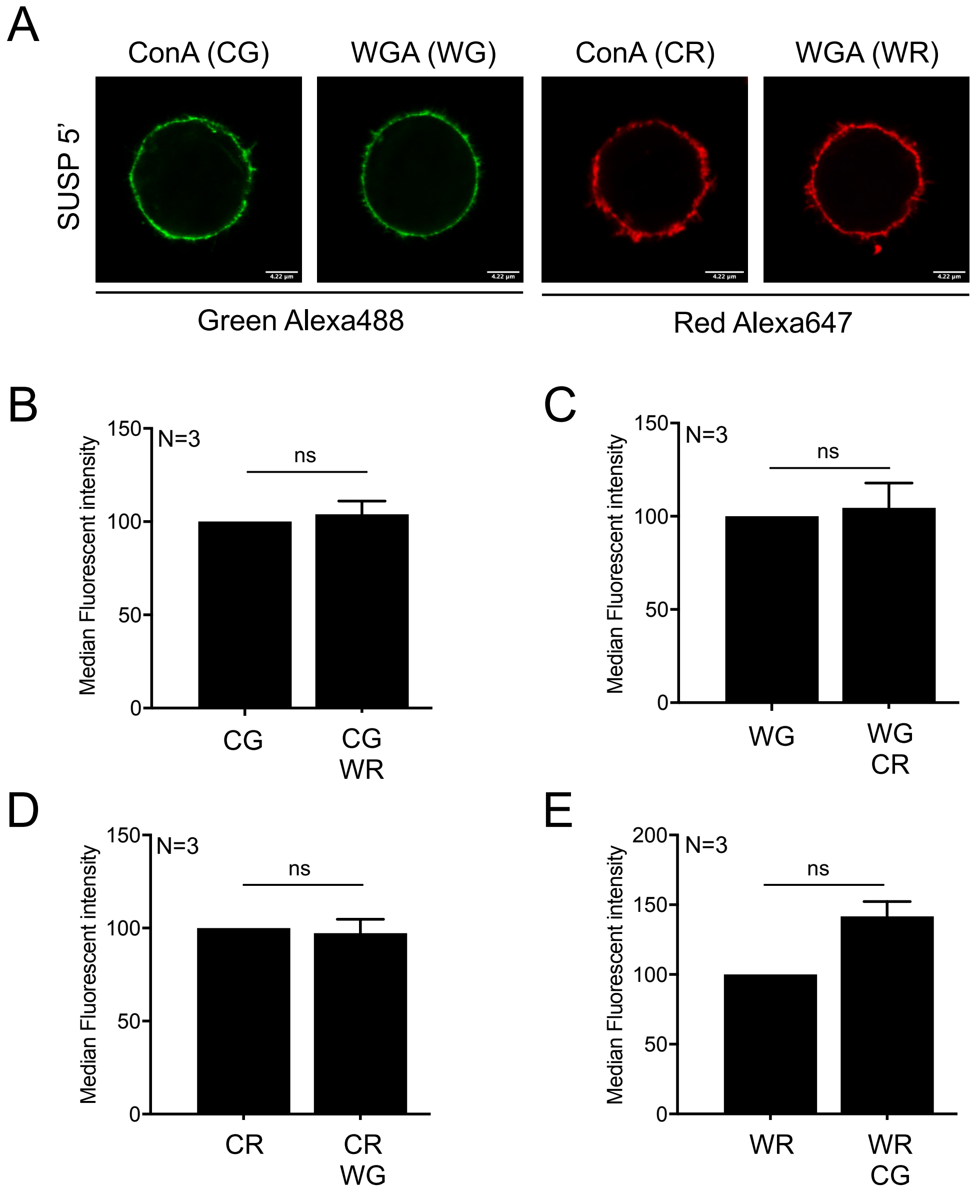


Fig. S1: Dual-labelling of cell surface glycans in non-adherent WT-MEFs with ConA and WGA lectins, conjugated to Alexa488 (green) or Alexa647 (red). (A) Confocal imaging data showing cell surface localization of lectins in non-adherent WT-MEFs. (B-E) Median fluorescence intensities from flow cytometry, for non-adherent WT-MEFs labeled with one lectin or with a pair of lectins conjugated to different fluorophores (CG, ConA-Green; WG, WGA-Green; CR– ConA-Red; WR, WGA-Red). Fluorescence values are normalized by the respective single-labeled controls. Statistical analysis was done using the two-sided single sample Wilcoxon $t$-test with a $p$-value cutoff of $.05$.

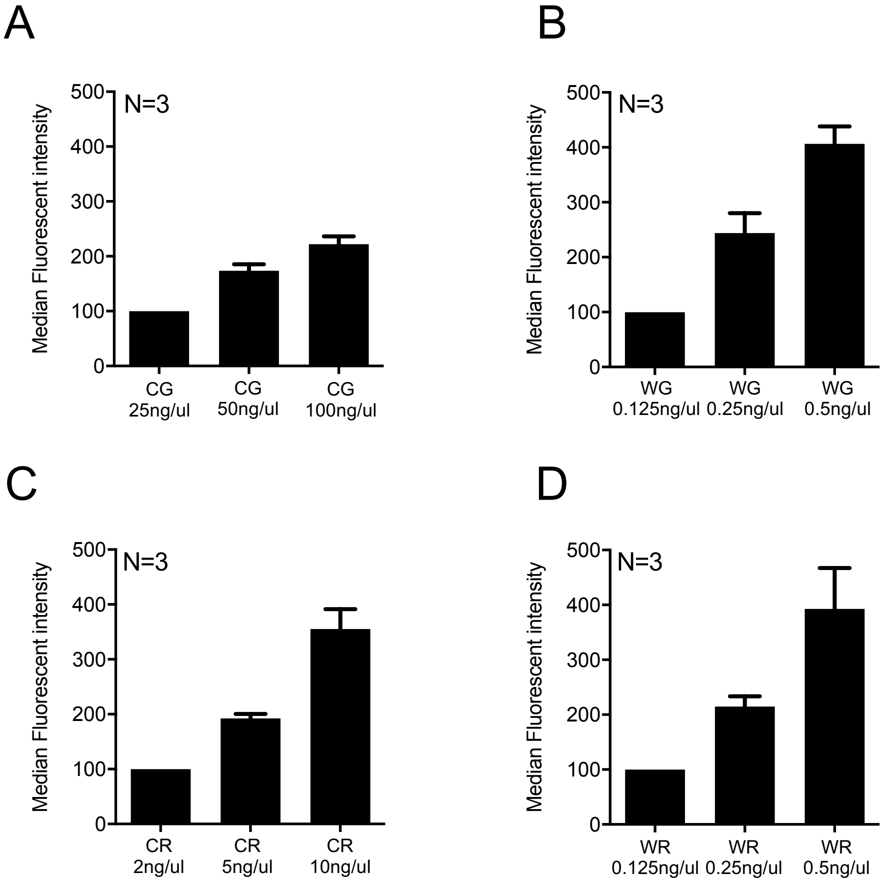


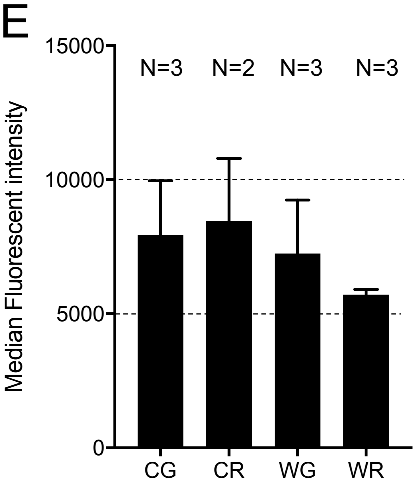


F

| Lectin | Binding specificity | Concentration |
|---|---|---|
| ConA Green | Mannose | 50ng/ul |
| ConA Red | | 5ng/ul |
| WGA Green | N-acetylglucosamine Sialic acid | 0.5ng/ul |
| WGA Red | | 0.25ng/ul |

Fig. S2: Standardization of lectin concentrations for labelling. (A-D) Non-adherent WT-MEFs are detached and labelled with the indicated lectins. We show median fluorescence intensities from flow cytometry, normalized by the intensity at the lowest lectin concentration (CG, ConA-Green; WG, WGA-Green; CR–ConA-Red; WR, WGA-Red). (E) Raw values for median intensities at the lectin concentrations selected for labelling lie within a range of 5000-10000 units. (F) Table showing selected concentrations for each lectin, used throughout in the study.

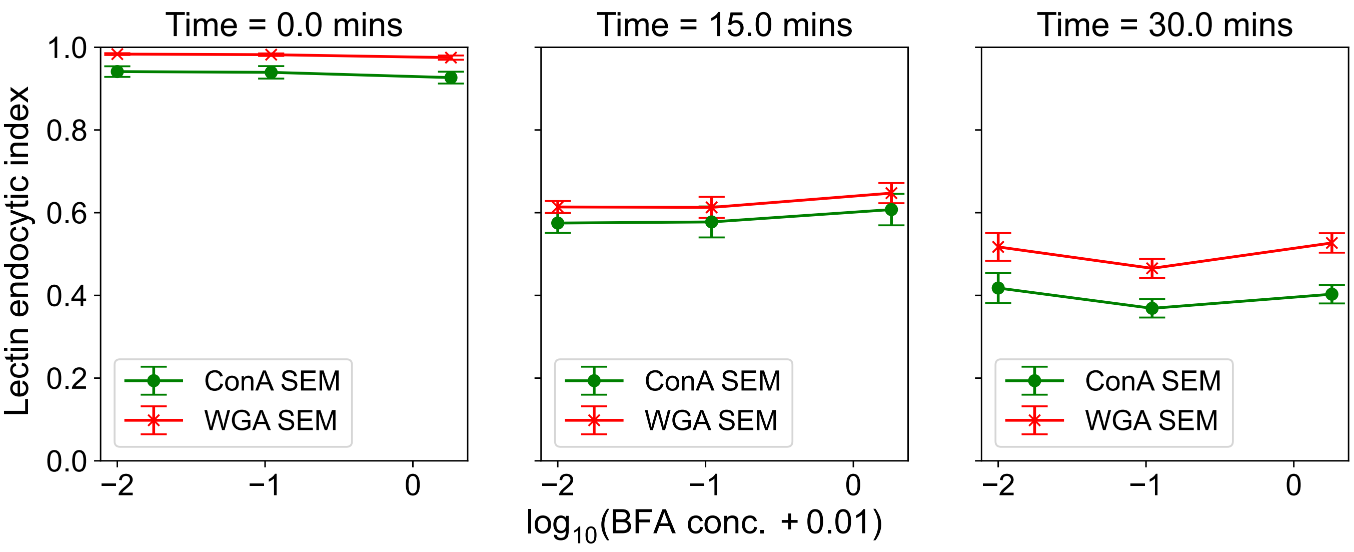


Fig. S3: Comparison of ConA and WGA endocytic indices across BFA concentrations at given times.

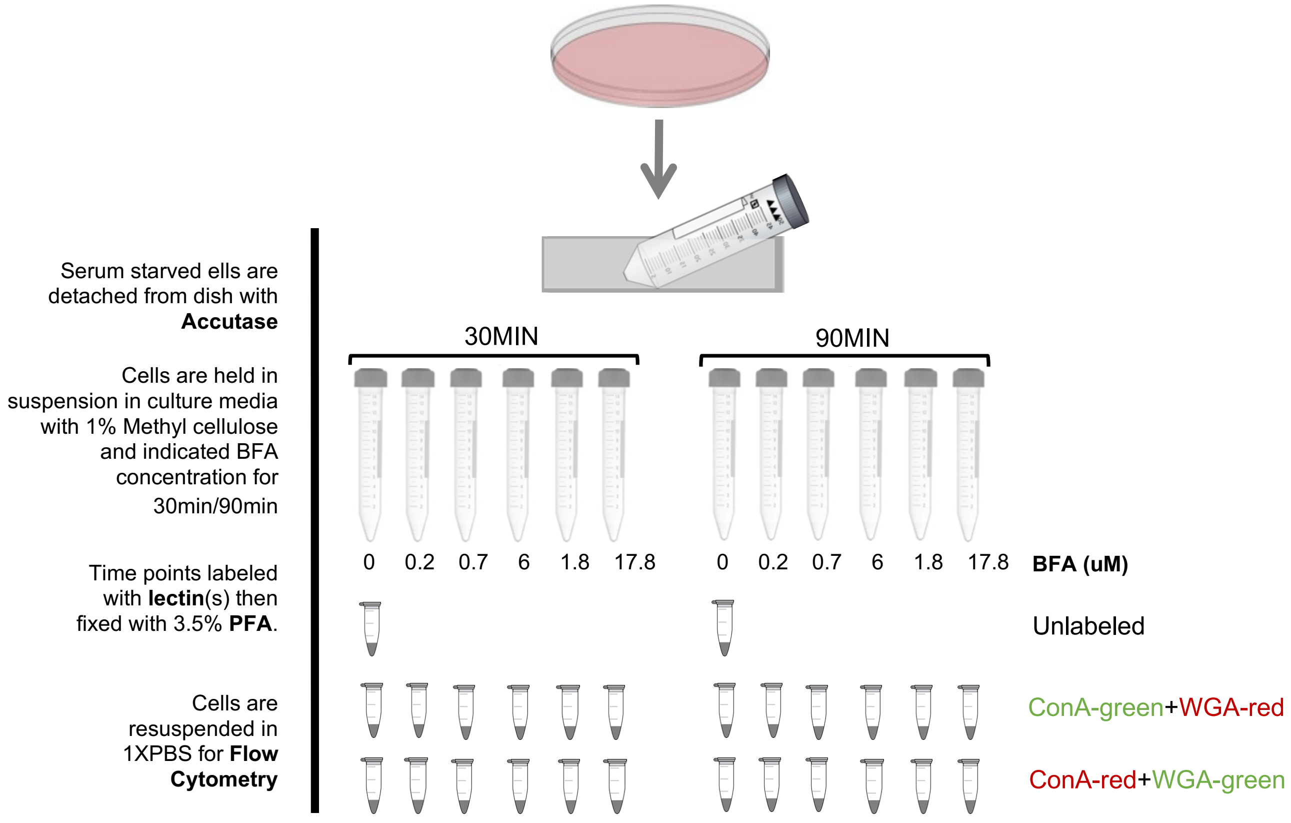


Fig. S4: Schematic of the lectin labelling experiment.

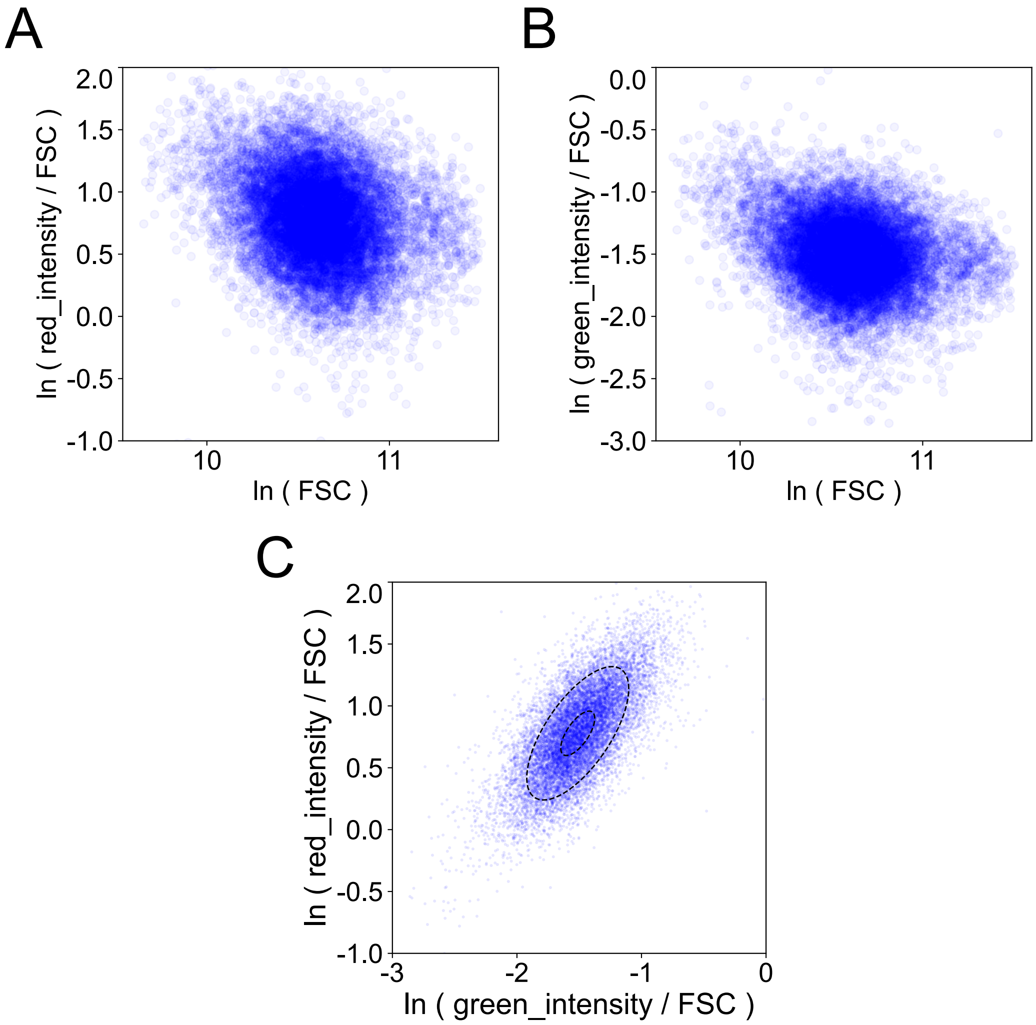


Fig. S5: Example of flow-cytometric single-cell lectin data at 0 μM BFA and 30 min, for ConA-Green / WGA-Red lectins ($n = 13,882$ cells). (A,B) Lectin signal intensities after scaling by forward scatter (FSC). (C) Scaled red vs. green intensities. We use robust estimation to find the parameters of the best-fit bivariate log-normal distribution. Ellipses show the $1\sigma$ and $3\sigma$ contours of the distribution.

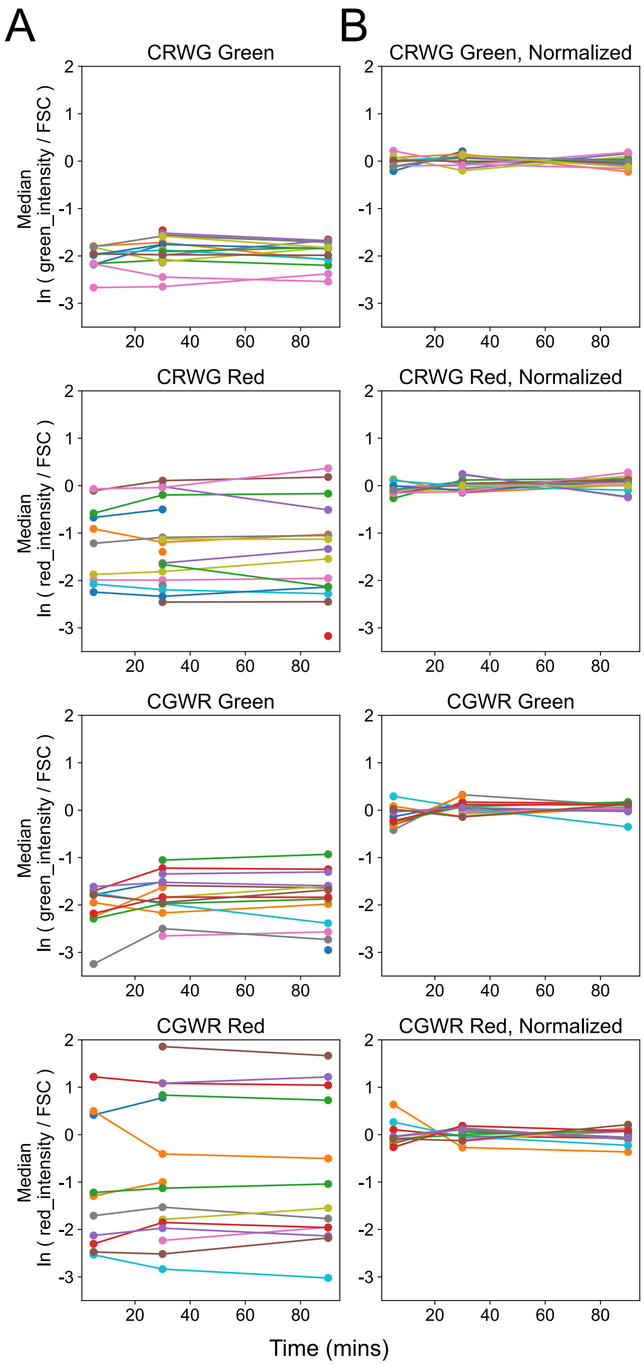


Fig. S6: Batch-to-batch normalization. We show median red and green intensities of cell populations at 0 μM BFA and various time points. (A) Raw intensities. (B) Intensities after normalizing by the geometric mean of the 5 min, 30 min and 90 min values. Batches were retained if the BFA 0 μM control was available for at least one time point.

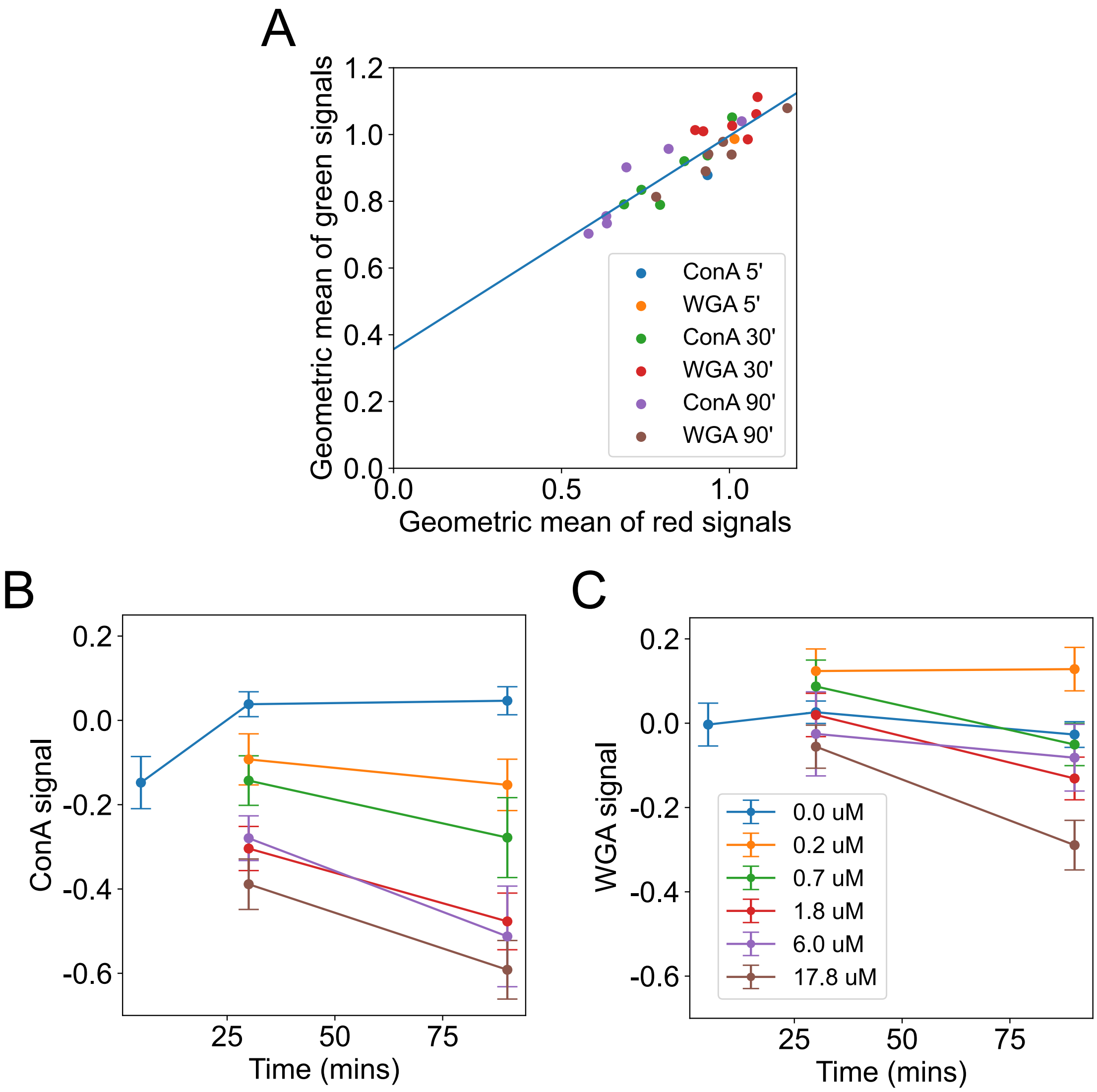


Fig. S7: (A) Comparison of fluorophore-swapped lectin measurements. Each point corresponds to a specific BFA concentration, time, and fluorophore configuration, and shows the geometric mean of the signal across replicates. (B,C) Dynamics of ConA and WGA signals. We convert CRWG experiments to CGWR units by using the linear relationship in Fig. S7A, then pool all replicates. Points show mean ± SEM across replicates in log space (natural logs). For 0 μM BFA the ConA signal (but not the WGA signal) transiently drops at the 5 min time before recovering. This may be due to reconfiguration of intra-Golgi trafficking upon cell de-adhesion.

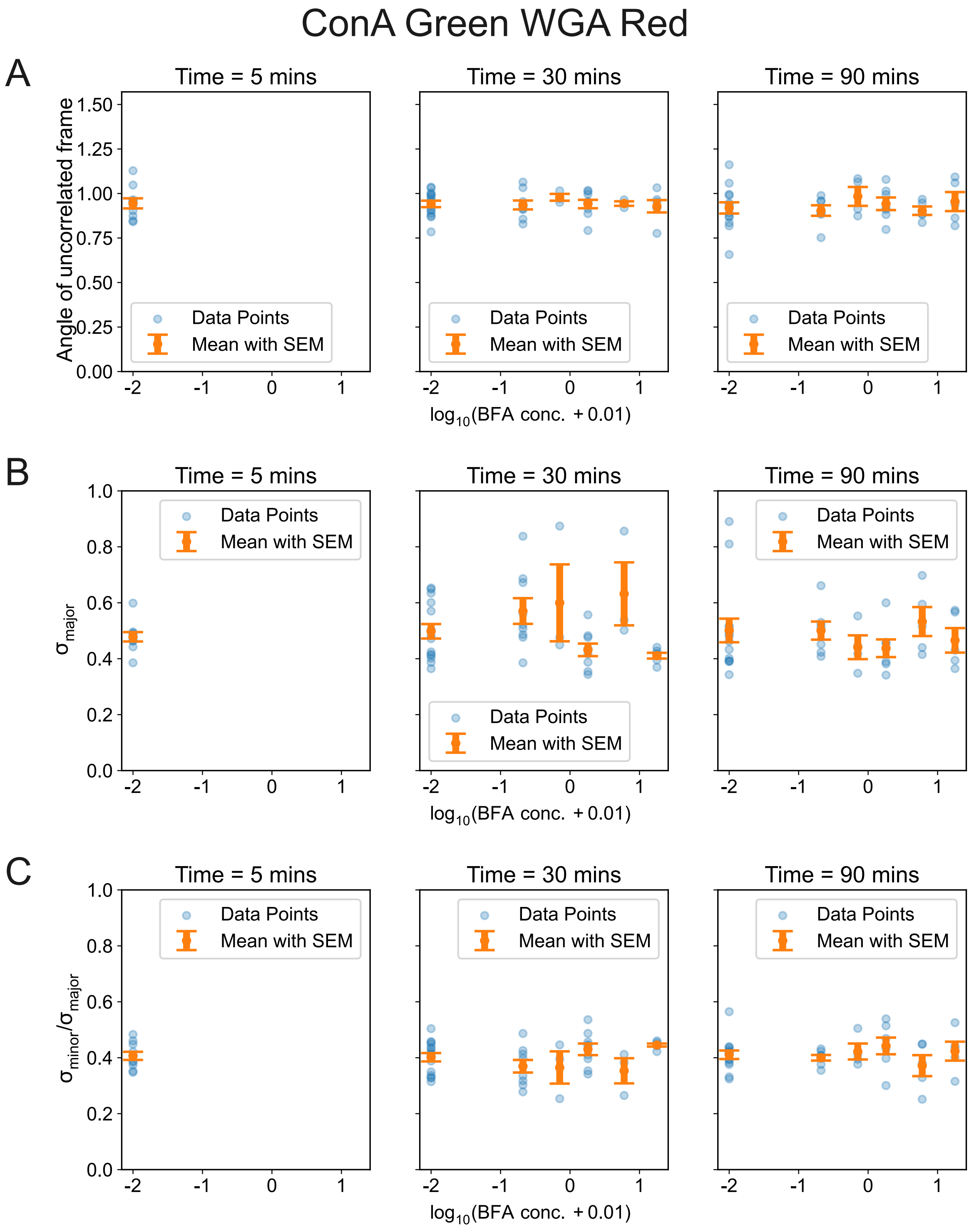


Fig. S8: Covariances of the ConA-Green WGA-Red lectins as a function of time and BFA concentration. We show parameters for the $1\sigma$ elliptical contours of log-normal distributions in log-log space (natural logs), with green signals on the X-axis and red signals on the Y-axis (Fig. S5C). (A) Angle in radians of the uncorrelated frame of the ellipse, with respect to X-axis. (B) Length of the major axis. (C) Ratio of the minor to major axes.

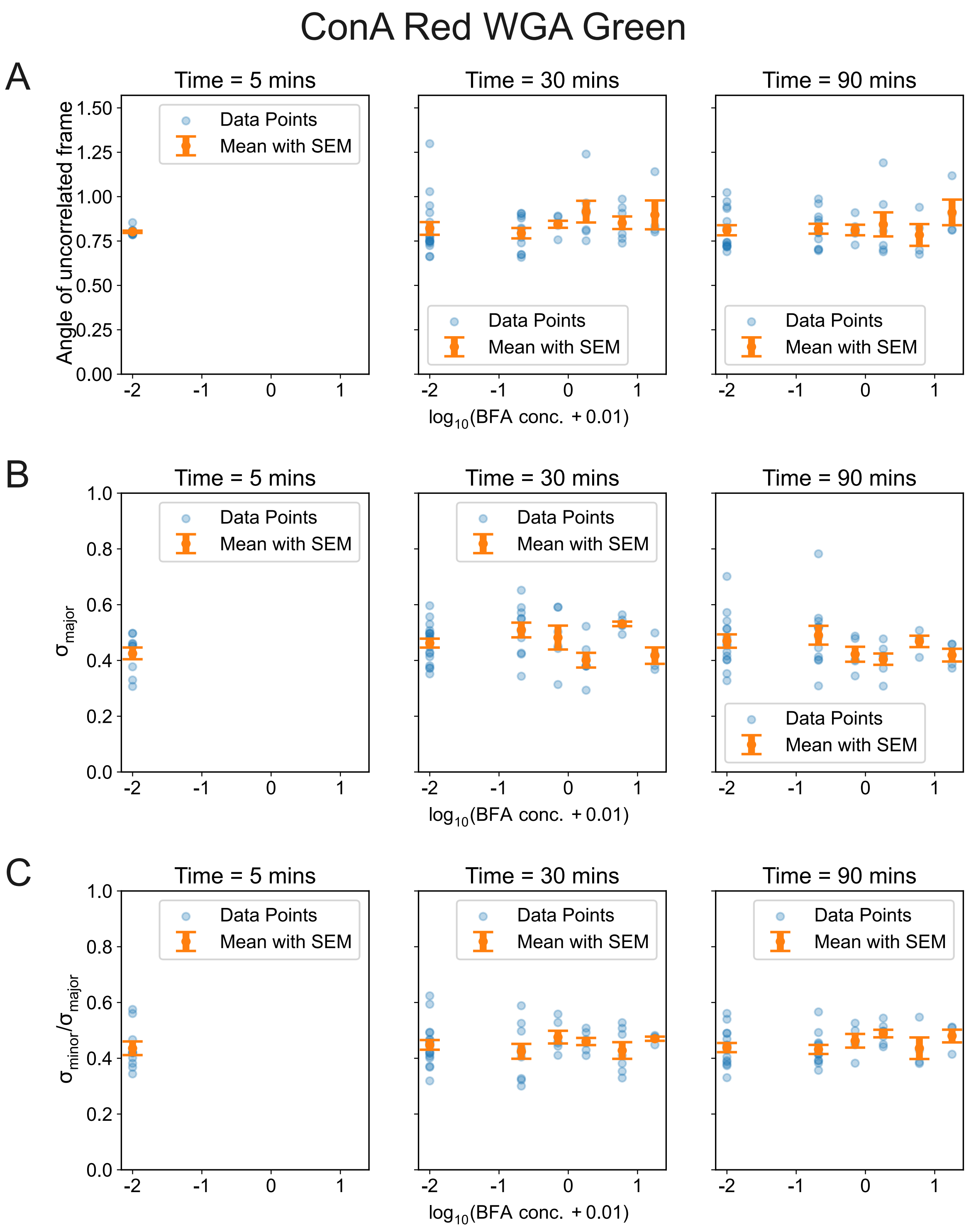


Fig. S9: Covariances of the ConA-Red WGA-Green lectins as a function of time and BFA concentration. We show parameters for the $1\sigma$ elliptical contours of log-normal distributions in log-log space (natural logs), with green signals on the X-axis and red signals on the Y-axis (Fig. S5C). (A) Angle in radians of the uncorrelated frame of the ellipse, with respect to X-axis. (B) Length of the major axis. (C) Ratio of the minor to major axes.

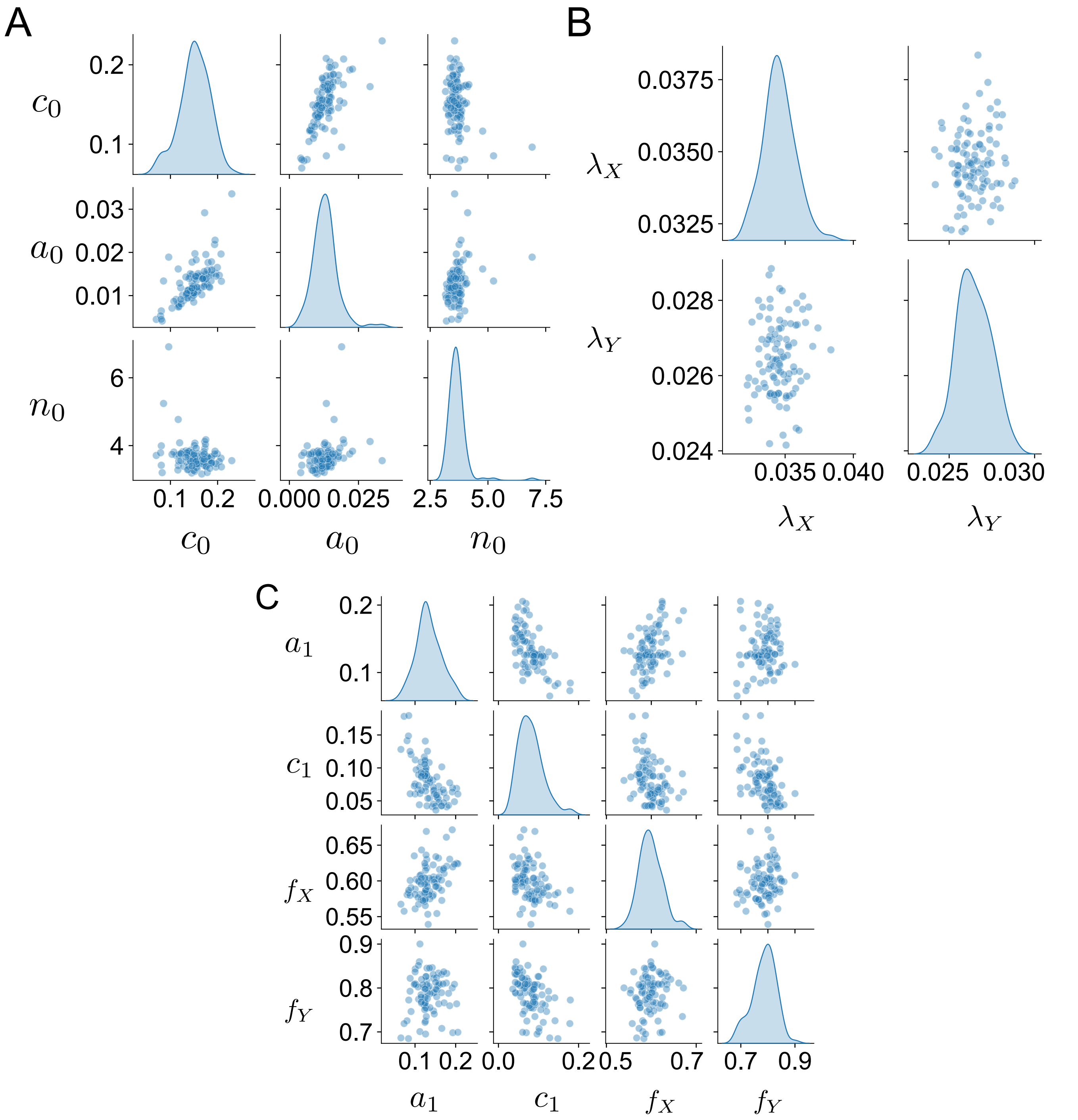


Fig. S10: Bootstrap estimates of fitted parameter uncertainties. We show fitted parameters from 100 bootstrap replicates for (A) the Golgi fragmentation model, (B) endocytosis rates, and (C) cell-surface glycan dynamics. For comparing with parameter values in Table 1, use: $a_0 = 1/t_0$, $a_1 = \gamma_0/\beta$, $f_X = R_X/(R_X + \alpha n_0)$, $f_Y = R_Y/(R_Y + \alpha n_0)$.

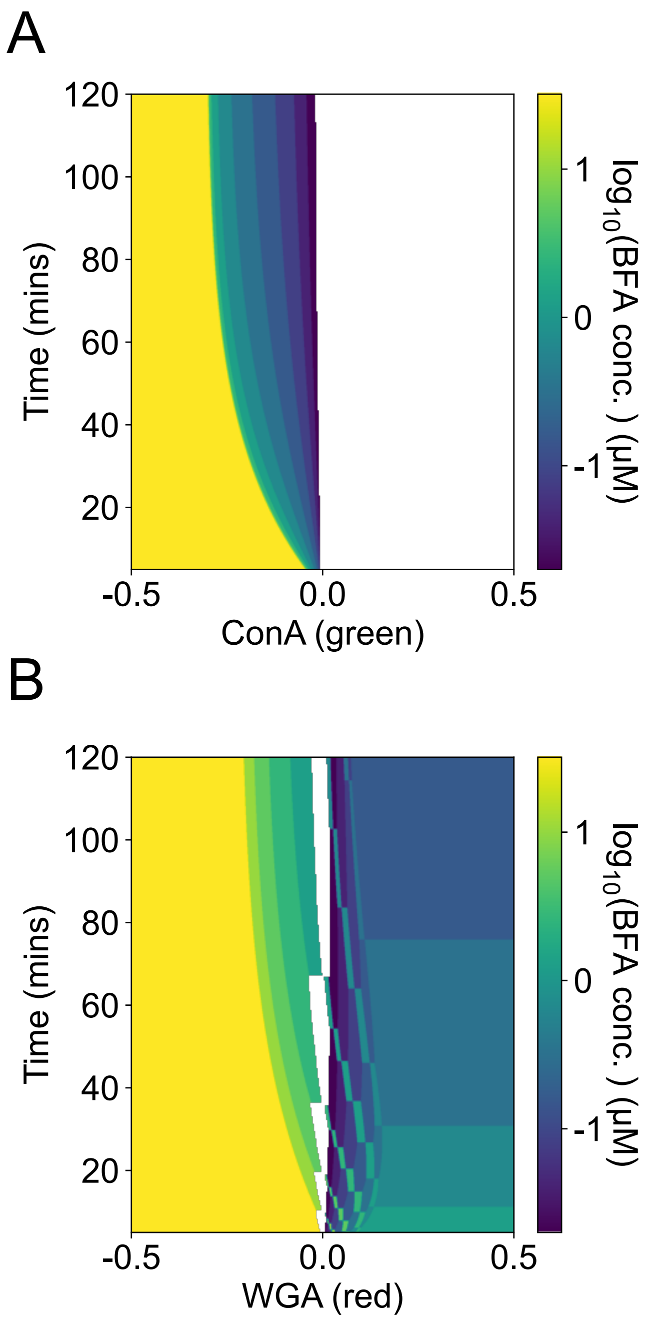


Fig. S11: Single-lectin decoders for $N = 12$ populations, across time. (A) Decoder using ConA alone. (B) Decoder using WGA alone. Colors show the best-guess BFA concentration value at each level of green or red. White represents 0 μM BFA.

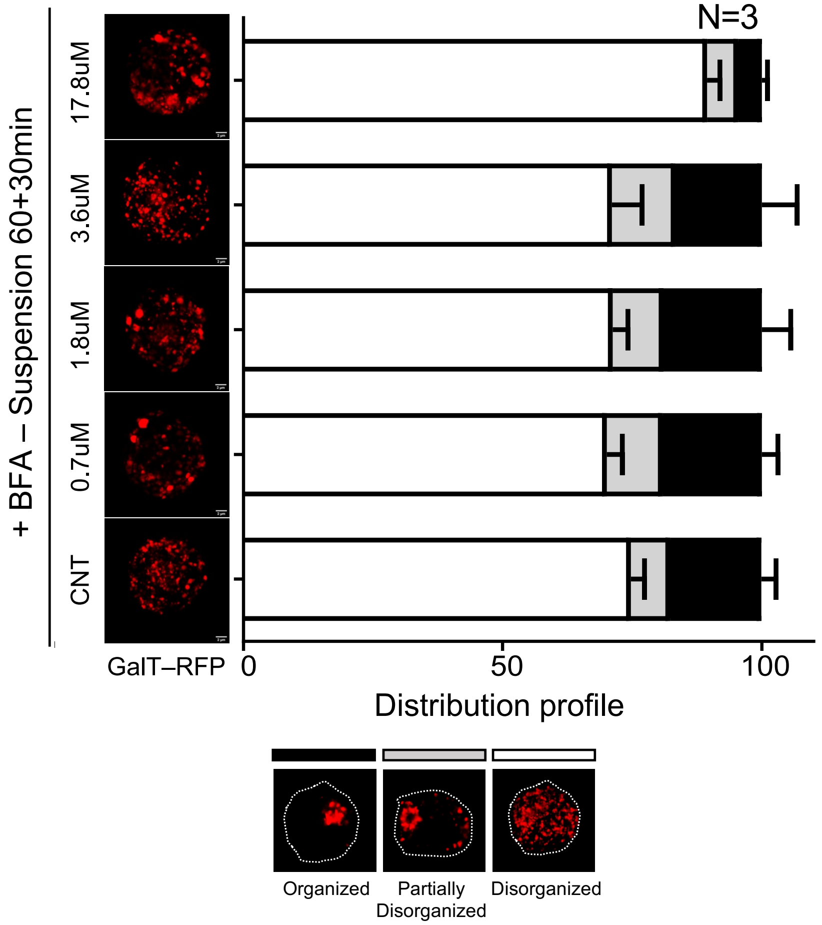


Fig. S12: BFA does not appear to impact trans-Golgi morphology. We image the trans-Golgi in non-adherent WT-MEF cells using a fluorescently labeled marker (GalTase-RFP). Bottom: representative images showing the three trans-Golgi morphology classes. Top: Distribution of trans-Golgi morphologies in cell populations ($n > 100$) suspended for 60 min, then incubated for 30 min with varying concentrations of BFA. Error bars show the SEM over $N$ replicate populations.
